# Arrested Agonist Paradigm For Selective Radiosensitization of Prostate Cancer

**DOI:** 10.1101/2023.10.01.560227

**Authors:** Jonathan B. Coulter, Michael C. Haffner, Yonggang Zhang, Haoming Zhou, Minh-Tam Pham, Jiayu Chen, Roshan Chikarmane, Alok Mishra, Maire S. Mehl, Stella Kazibwe, Kirsten Choi, Ava Archey, Sarayu Valluri, Shawn E. Lupold, Daniel Song, Angelo De Marzo, William G. Nelson, Theodore L. DeWeese, Srinivasan Yegnasubramanian

## Abstract

As a prototypical nuclear hormone receptor, the androgen receptor (AR) signals via a sequential cascade triggered by binding to androgenic ligands such as testosterone and dihydrotestosterone (DHT). This cascade includes dimerization of the ligand-receptor complex, nuclear translocation, chromatin binding to response elements, recruitment of TOP2B and co-activator complexes, and induction of an effector transcriptional program. In prostate cancers, this AR signaling cascade is an essential driver of growth and survival, yet its activity confers potential vulnerabilities through transient TOP2B-mediated DNA double strand breaks. We investigated the ability of non-steroidal AR ligands to activate initial steps of the AR signaling cascade up to the point of AR- and TOP2B-mediated double strand breaks, with subsequent arrest of the signaling cascade to prevent induction of pro-growth/survival transcriptional programs in prostate cancer cells. We identified hydroxyflutamide (FLU) as such an androgen receptor arrested agonist; in androgen-deprived conditions, FLU induced AR nuclear translocation, chromatin binding, and TOP2B-mediated double strand breaks, but failed to induce AR target gene expression and prostate cancer cell growth. The FLU-mediated arrest in the signaling cascade could be attributed to the inability of FLU to allow association of AR with SMARCD2, a critical component of the BAF chromatin remodeling complex required for androgen induced AR co-activation. Interestingly, the FLU-induced, AR- and TOP2B-mediated double strand breaks could be used to selectively sensitize AR-positive prostate cancer cells to ionizing radiation *in vitro* and *in vivo*. These findings support a novel arrested agonist paradigm for selective radiosensitization of prostate cancer cells without inducing AR-mediated pro-growth and survival transcriptional programs.

## Introduction

The demonstration of androgen dependence in prostate cancer in 1941 provided the underlying basis for anti-androgen therapies in prostate cancer treatment^1^, which remain central to treating both locally-advanced and metastatic disease. The dependence on androgen receptor (AR) signaling in prostate cancer cells is remarkable as identifying mechanisms of circumventing androgen blockade remains an active area of investigation, and novel AR antagonists continue to provide survival benefit in treating metastatic prostate cancer even in patients on androgen deprivation therapies^2-5^. The AR and other nuclear hormone receptors bind their respective ligands and mediate transcription, differentiation, and growth through response elements (RE) in various genomic sequences. In the absence of ligand, the AR is sequestered in the cytoplasm and bound to heat-shock proteins (HSPs). Upon androgen-binding, conformational changes in the AR promote its dissociation from HSPs and translocation to the nucleus, where it dimerizes and binds as a complex with coactivators, general transcription factors, RNA polymerase II, and other proteins at AREs^6,7^. This complex mediates subsequent activation of genes directly. The program of genes transcribed downstream of AR signaling are known to be involved in numerous functions, such as cellular differentiation and secretion of components of semen, the most well-known of which is KLK3 encoding PSA which is commonly measured to inform the growth of prostate cancer, metastasis following treatment, and response to treatments^6^.

Interestingly, bicalutamide, a clinically used non-steroidal AR antagonist, has been shown to induce AR nuclear translocation and DNA binding in the absence of androgens, yet the AR forms a transcriptionally inactive complex with respect to initiating transcription and prostate cell growth^8^. Additionally, flutamide, a first generation AR antagonist has weak agonist activity in wildtype AR and stronger agonist activity even driving cell growth in the context of specific AR mutations such as the T877A mutation in the ligand binding domain of the AR identified in the LNCaP cell line^9-11^. A prevailing explanation of the pro- or anti-transcription effects of various AR ligands involves ligand-specific changes in AR conformation resulting in differential recruitment of co-factors necessary for transcription to induce downstream cell growth in prostate cancer cells^12^. The myriad context-specific factors resulting in full-agonist or full-antagonist activity suggest the possibility that some ligands in specific contexts may act as “arrested agonists,” inducing AR conformations resulting in an arrest throughout various points along the AR signaling pathway. Such arrested agonists could potentially induce initial steps of the AR signaling cascade yet induce a conformational change that fails to promote assembly of key transcription coactivators.

Reports demonstrating that AR signaling causes transient double strand breaks (DSB)^13^ and radiosensitizes AR-expressing prostate cancer cells^14^ suggest vulnerabilities reside downstream of AR ligand-binding despite its importance in driving prostate cancer growth. It may therefore be possible to exploit such vulnerabilities using the putative arrested agonist activity of some AR ligands for therapeutic benefit. Here we investigated the androgen receptor as a model of the prototypical nuclear hormone system. In a screen of FDA-approved non-steroidal anti-androgens, we identified hydroxyflutamide (OH-FLU), the active metabolite of the clinically used anti-androgen flutamide, as an AR ligand with arrested agonist activity. We demonstrate that OH-FLU differentially recruits coactivators compared to the natural ligand dihydrotestosterone (DHT). While OH-FLU induced AR nuclear translocation, chromatin-binding, and DNA damage, leading to significant sensitization to ionizing radiation, pro-growth and survival pathways were not activated. The topological constraints encountered by AR complexes are likely relieved at least in part by topoisomerase 2B (TOP2B), which is known to play a role in DSB formation induced by AR signaling^13^. We therefore suggest that ligands such as OH-FLU be considered arrested agonists with potential to exploit DNA damage induced by nuclear hormone receptor signaling pathways while avoiding pro-growth and survival pathways in the treatment of tumor cells which rely on them for growth and survival.

## Results

In order to characterize the various, potentially unique programs along the AR cascade in response to various ligands, we first evaluated AR subcellular localization in response to FDA approved, clinically used AR antagonists in the background of castrate levels of androgens. Androgen-deprived LAPC4 cells were exposed to 10μM of various clinically-used, non-steroidal AR antagonists alone for 2 hours or 10nM dihydrotestosterone (DHT) to screen for their ability to induce AR nuclear translocation. Hydroxyflutamide (OH-FLU) and bicalutamide (BIC) exposure resulted in significant nuclear accumulation of AR, whereas enzalutamide (Enz), apalutamide (APA), nilutamide (NILU), and darolutamide (DARO) failed to induce significant nuclear localization of AR compared to vehicle control (**Fig. 1A, 1B, S1**). While all AR antagonists have been shown to prevent full AR-mediated signaling in the context of competing with AR agonists, nuclear AR translocation in the absence of agonists has been demonstrated by BIC and FLU, resulting in an inactive complex at known AR binding sites^7^. In androgen-deprived conditions, FLU and BIC, but not ENZ or APA, induced AR binding at *KLK3* and *TMPRSS2* enhancers (**Fig. 1C**), consistent with previous reports. While FLU and BIC are known to induce AR nuclear translocation and binding to AR response elements (ARE), it is possible that their activities induce gene expression of a canonical or non-canonical AR-mediated transcription program. Neither FLU, BIC, ENZ, nor APA led to substantial induction of gene expression compared to DHT exposure (**Fig 1D, 1E, and S2**), suggesting that binding of AR induced by FLU or BIC resulted in inactive AR complexes at AREs. To further characterize the effects of these AR-interacting compounds in the AR signaling pathway, cell growth in castrate conditions following a single stimulation of AR antagonists was measured, where no antagonist led to significant cell growth compared to vehicle control (**Fig. 1F, S3**). While no AR antagonist promoted canonical AR target gene expression or cell growth in the background of castrate levels of androgen, individual antagonists displayed inhibition of the AR signaling pathway at different points along AR signaling steps (**Fig. 1G**).

**Figure 1:**
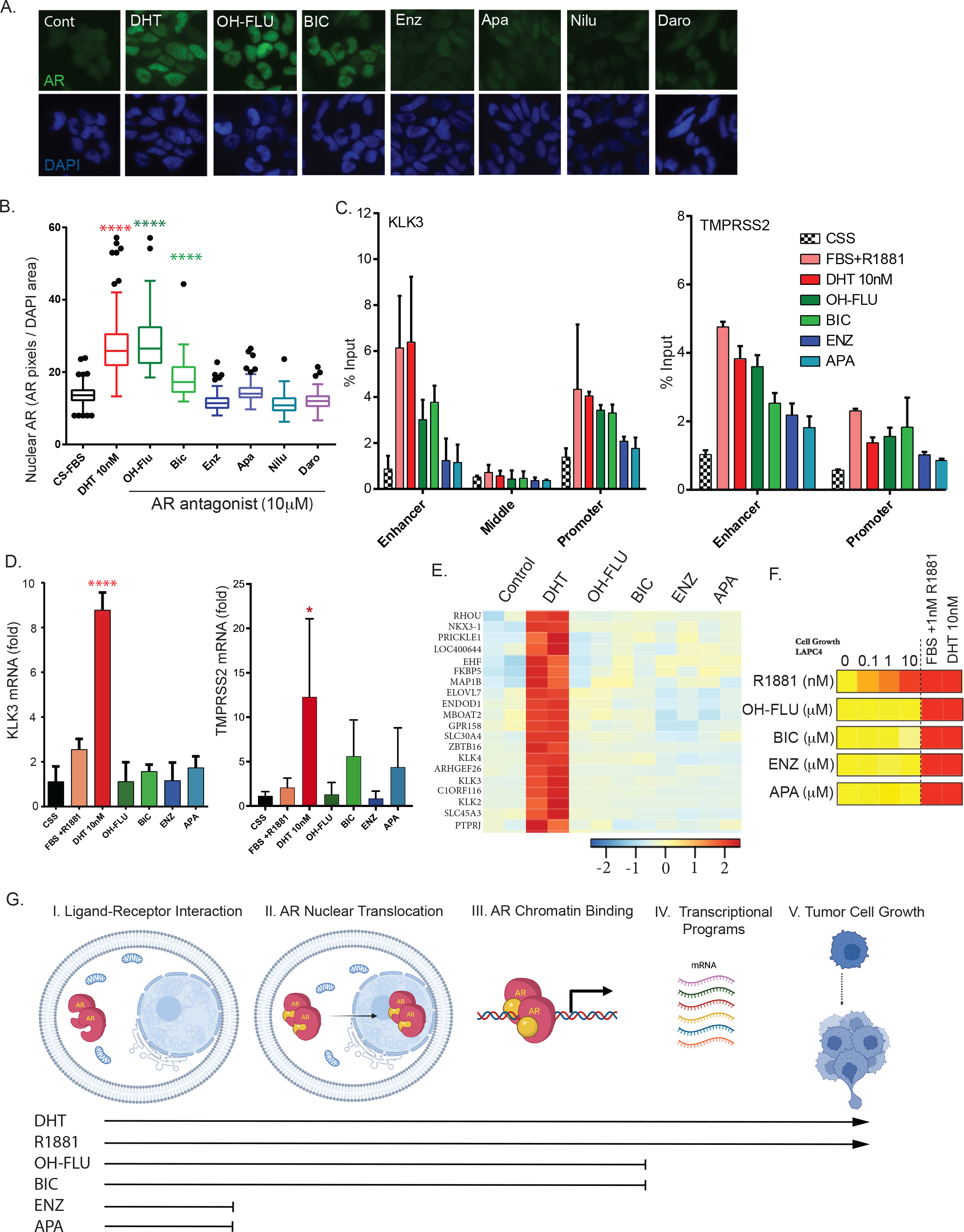
Androgen receptor ligands show complete agonist, complete antagonist, and arrested agonist activity. **(A)** Representative immunofluorescent detection and **(B)** quantification of AR (green) and DAPI (blue) in LAPC4 cells stimulated for 2h with vehicle (Cont), 10nM DHT, or 10mM hydroxyflutamide, bicalutamide, enzalutamide, apalutamide, nilutamide, or darolutamide. (n=3) **(C)** AR binding at the KLK3 ARE-containing enhancer and promoter and ARE-lacking middle region (left) and TMPRSS2 enhancer and promoter (right) following stimulation of androgen-deprived LAPC4 cells with the indicated AR ligand (antagonists 10mM) for 6h. Values shown are % of input DNA (n=3). **(D)** qRT-PCR measurement of KLK3 (left) or TMPRSS2 (right) mRNA in androgen-deprived LAPC4 cells stimulated with indicated AR ligand. Values are fold change over vehicle-treated (CSS) state and TBP was used as a reference gene. ****p<0.0001 (n=3), *p<0.05 **(E)** Gene expression changes following stimulation of androgen-deprived LAPC4 cells with DHT (10nM) or indicated antagonist (10mM) measured by RNAseq. Heatmap shows top 20 genes changed following DHT exposure (n=2 biological replicates). **(F)** Heatmap of cell growth by incucyte of androgen-deprived LAPC4 cells following a single stimulation with indicated AR ligand (10mM) over 7 days. **(G)** Scheme depicting Arrested Agonist concept. Lines indicate ability for indicated AR ligand to promote AR signaling steps. *Abbreviations: AR, androgen receptor; DHT, dihydrotestosterone; OH-FLU, hydroxyflutamide; BIC, bicalutamide; ENZ, enzalutamide; APA, apalutamide*.

The observation that an AR antagonist could induce AR nuclear translocation and chromatin binding yet fail to induce gene transcription or cell growth as full agonists, led us to hypothesize that AR conformation changes may be induced in a ligand-specific manner, leading to formation of differential AR-interactomes. In order to identify ligand-specific AR interactions following brief agonist or antagonist exposures, endogenous AR-RIME was conducted in LAPC4 cells in the background of androgen deprivation following 16 hours of exposure to vehicle, growth medium, DHT, or FLU. The AR-interacting proteins observed following agonist exposure included expected AR coactivators such as FOXA1, HOXB13, and EP300, as well as nearly all members of the canonical BAF Complex (cBAF) (**Fig 2A**). The most abundant AR-interacting proteins in DHT-treated cells compared to OH-FLU-treated cells included cBAF members, suggesting cBAF-AR interactions are induced preferentially in DHT-treated cells compared to OH-FLU treated cells (**Fig 2B, 2C**). AR-RIME conducted at 3 hours following stimulation of androgen-deprived cells with DHT or OH-FLU revealed 42 AR-associated proteins, of which members of the nucleosome remodeling Brg/BRM Associated Factor (BAF) (SWI/SNF) complex were highly represented. Of note, the BAF complex member SMARCD2 was found to be the most highly abundant AR interactor at this timepoint following stimulation with DHT (**Fig 2B, 2C**). The finding that SMARCD2 was identified as the most abundant DHT-induced BAF complex member at both 3 and 16 hour timepoints, and SMARCD2 was the most abundant AR-interacting protein overall at the 3 hour timepoint, suggests the possibility that this BAF complex member may represent an early and enduring coactivator of AR signaling through AR agonist exposure. Reciprocal co-immunoprecipitation revealed that AR and SMARCD2 interactions occurred only in the context of DHT stimulation (**Fig 2D**). Similarly, proximity ligation assays measuring AR-SMARCD2 interactions demonstrated that DHT, but not FLU, induced interaction of AR and SMARCD2 in LAPC4 and VCaP (**Fig 2E**). Interestingly, while no antagonist led to an increase in growth in LAPC4 cells (**Fig S3**), both DHT and FLU exposure led to AR-SMARCD2 interaction in LNCaP, a cell line known to grow in the presence of FLU in the background of castration (**Fig S1**), an effect attributed to a T877A mutation in the AR ligand binding domain conveying an antagonist-to-agonist switch^9^.

**Figure 2:**
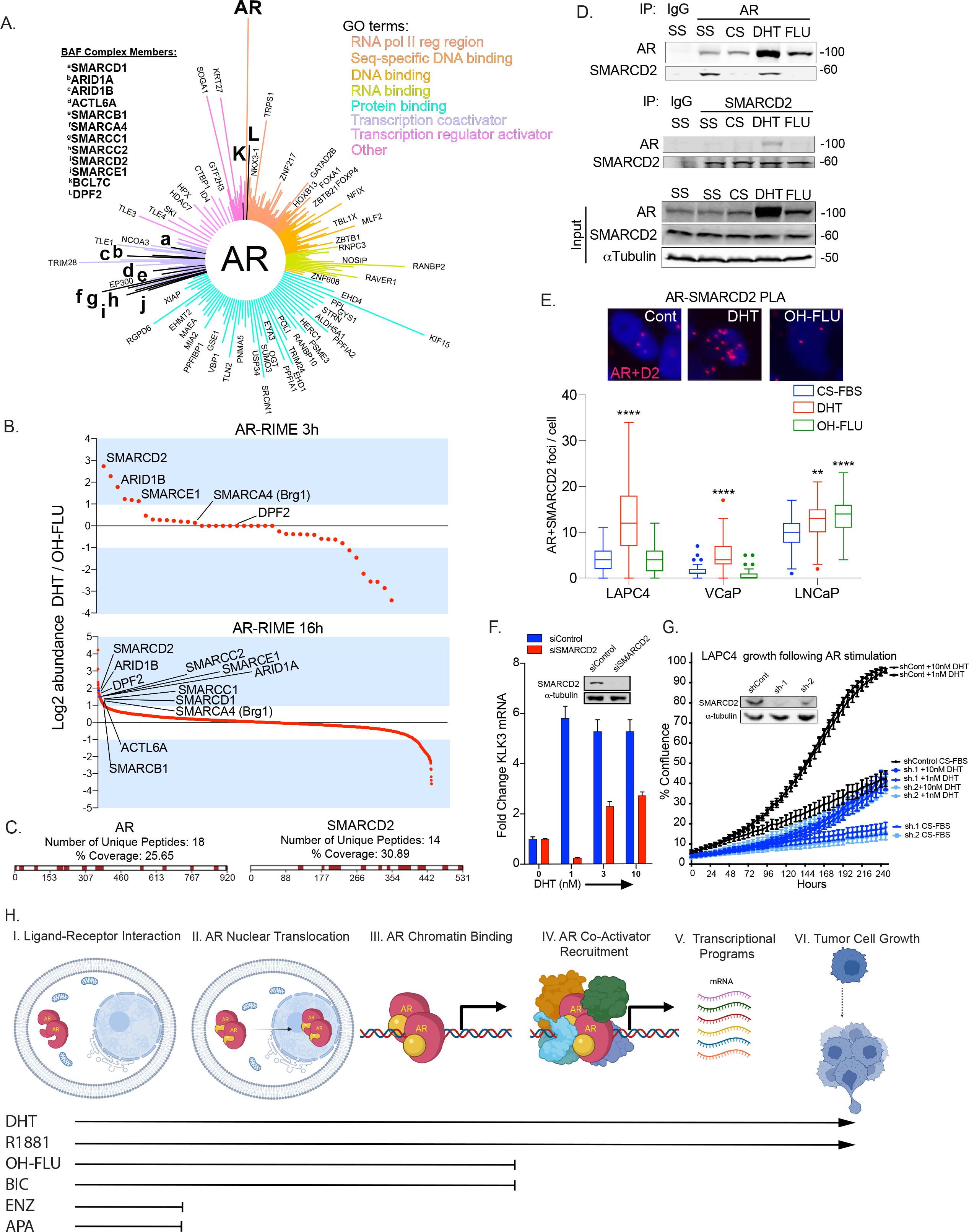
Differential BAF complex recruitment by complete and arrested AR agonist stimulation. **(A)** MS-ARC plot of AR-RIME results. Proteins identified following non-immune IgG have been removed, and AR-associated proteins of all conditions are clustered according to color-coded GO-term molecular functions (top right of Panel A). Line length from the center represents the Mascot score for the given protein. Bolded lines labeled A-L are members of the cBAF complex (labels are shown in top left) **(B)** Waterfall plot of AR-RIME results showing AR interactions in LAPC4 cells stimulated with DHT (10nM) or OH-FLU (10μM). Proteins shown represent ratios of DHT- to OH-FLU-induced interactions at 3 hours (top) or 16 hours (bottom). Coverage of AR and SMARCD2 are shown in **(C). (D)** Representative co-immunoprecipitation of AR (top) and SMARCD2 (bottom) following stimulation of steady-state androgen levels (SS), androgen-deprived LAPC4 cells (CS), or stimulated with DHT or OH-FLU for 16 hours. Input levels of AR, SMARCD2, or a-tubulin are shown below. **(E)** Proximity Ligation Assay results for AR and SMARCD2 interaction. Representative images (top) showing interaction foci and foci quantification via Telometer demonstrated in box-and-whiskers plot following stimulation of androgen-deprived LAPC4, VCaP, and LNCaP cells with 10nM DHT or 10μM OH-FLU. Statistical significance was determined by 1-way ANOVA and Dunnett post-test compared to androgen-deprived condition. **p<0.01, ****p<0.0001 **(F)** KLK3 mRNA was measured by qRT-PCR. Knockdown of SMARCD2 using siRNA led to a decrease in KLK3 levels in cells deprived of androgens and stimulated with DHT for 24 hours. **(G)** Growth of control (shEV) or SMARCD2 knockdown (sh.1 or sh.2) measured using incucyte. LAPC4 cells were imaged at 6 hour intervals in androgen-deprived conditions (CS-FBS) or following a single stimulation with vehicle or the indicated concentration of DHT. Western blot shows SMARCD2 and a-tubulin in LAPC4 cells transduced with lentiviral particles containing an empty vector (EV) or 1 of 2 shRNAs targeting SMARCD2 (sh.1 or sh.2). **(H)** Scheme depicting Arrested Agonist concept. Note AR coactivator recruitment step appears to be a divergent step in AR ligand specificity comparing a full agonist (DHT) with a putative Arrested Agonist (OH-FLU or BIC).

The finding that SMARCD2 associated with AR only under conditions where cell growth was observed presented the notion that SMARCD2 could be a necessary coactivator for full AR program activation. To test this, KLK3 mRNA was measured following DHT stimulation with control or SMARCD2 knockdown. Knockdown of SMARCD2 led to a decrease in DHT-induced KLK3 and KLK2 mRNA in both LAPC4 (**Fig 2F, S4**) and VCaP cells (**Fig S4**), suggesting that SMARCD2 is required for efficient AR signaling. To determine the requirement for SMARCD2 in AR-mediated prostate cancer cell growth, LAPC4 cells transduced with an control or one of two shRNAs targeting SMARCD2 were plated in androgen-replete or deprived conditions and growth kinetics were measured. Knockdown of SMARCD2 led to a decrease in growth both in typical hormone-containing conditions (**Fig S4**) and in androgen-deprived cells given a single dose of DHT (**Fig 2G**), demonstrating that SMARCD2 is necessary for efficient AR-induced growth and further suggesting that AR ligands differentially promote coactivator recruitment required for full AR signaling (**Fig 2H**).

Given previous reports of AR agonist-induced recombinogenic^13^ and radiosensitizing^14^ DNA double strand breaks (DSB) we investigated the ability of AR antagonists to induce DSB while lacking the ability to initiate transcription and growth programs. In castrate conditions, 6 hour exposures to DHT or FLU led to increases in γH2A.x foci **(Fig 3A**) and frank DSBs as measured by the neutral comet assay (**Fig 3B**), while ENZ, which does not induce AR nuclear translocation, had no effect on DNA damage measures. As AR-mediated DSBs have been shown to involve topoisomerase 2B (TOP2B) activity^13^, we tested the requirement of TOP2B for FLU-induced DSBs in castrate conditions. Knockdown of either AR or TOP2B abrogated DSBs following DHT or FLU exposure, indicating a role for TOP2B in AR antagonist-induced DSB (**Fig 3C**), consistent with observations of TOP2B-mediated DSBs in AR agonist signaling. These results further dissect the AR signaling cascade and suggest that some AR antagonists such as FLU initiate an AR signaling cascade including initiating TOP2B-mediated DSBs yet fail to promote transcription through incomplete assembly of coactivators such as SMARCD2. We propose that the ligand-specific induction of DSBs as a potentially terminating step in the AR signaling cascade can be considered an “arrested agonist” endpoint which could be exploited for various purposes (**Fig 3D**).

**Figure 3:**
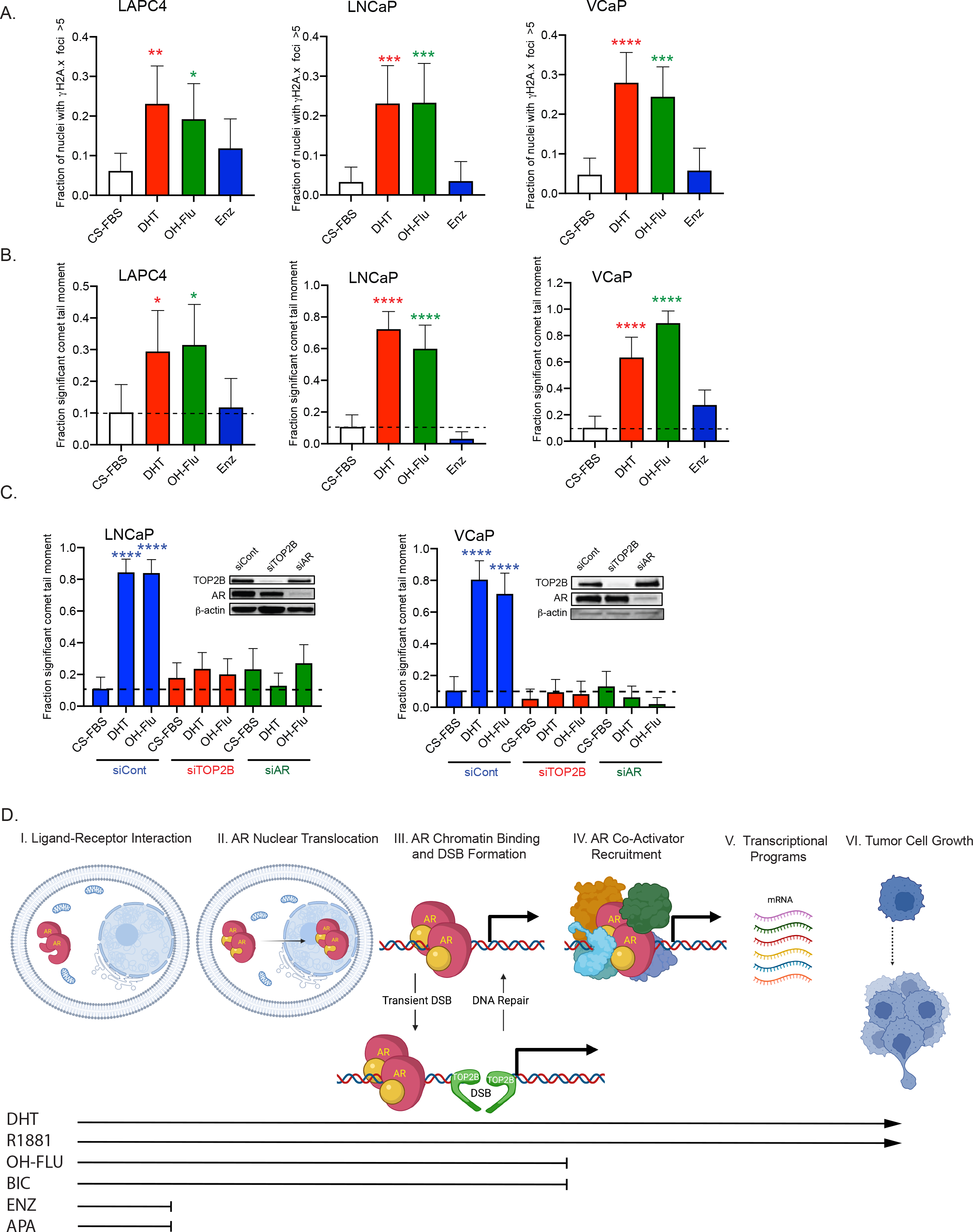
DNA double strand breaks induced by complete and arrested AR agonist stimulation. **(A)** Androgen-deprived LAPC4, LNCaP, or VCaP cells were stimulated with 10nM DHT, 10mM hydroxyflutamide, or enzalutamide for 6 hours and stained for γH2A.x and DAPI was used as a nuclear stain. Foci were counted using Telometer and bars indicate mean fraction of nuclei with > 5 foci. At least 50 nuclei were counted per experiment (n=3 experiments). **(B)** Androgen-deprived LAPC4, LNCaP, and VCaP cells were stimulated as described in A and at the end of 6 hours were placed directly into the neutral comet assay. The 90^th^ percentile of comet tail moment was determined for cells grown in androgen-deprived conditions and significance is defined as being above this threshold for all treatment groups. Results shown for at least 50 nucleoids per experiment (n=3 experiments). **(C)** siRNA against TOP2B or AR was transfected into LNCaP or VCaP cells prior to androgen deprivation and cells were stimulated with 10nM DHT or 10mM hydroxyflutamide for 6 hours and then placed into the neutral comet assay (n=2 experiments). *p<0.05, **p<0.01, ***p<0.001, ****p<0.0001 by 1-way ANOVA and Dunnett post-test. **(D)** Scheme depicting Arrested Agonist concept. Note AR Arrested Agonists (OH-FLU or BIC) induce AR- and TOP2B-mediated DSBs in the absence of significant AR target gene induction or promotion of tumor cell growth. *Abbreviations: DSB, DNA double strand break; TOP2B, topoisomerase-2-beta*.

In light of the promising preclinical^14^ and clinical^15^ data regarding use of AR full-agonists as selective DNA damaging therapeutic agents, our data suggest that FLU and other arrested agonists could be used as radiosensitizing agents which lack pro-growth effects. We therefore asked whether FLU-induced DSBs initiated immediately prior to the application of ionizing radiation (IR) resulted in synergistic DSB formation in cells with relatively low AR levels and lacking mutations in the ligand binding domain of the AR. Androgen-deprived LAPC4 cells were stimulated with various doses of FLU or DHT for 6 hours prior to doses of IR for immediate analysis by the neutral comet assay to measure DSBs. At relatively low doses of IR (0.5 and 1Gy), FLU pre-exposure resulted in formal synergy of DSB formation (**Fig 4A-B**), yet this synergy was absent as radiation dose increased, suggesting the DSBs observed were driven by IR at these doses. To determine whether synergistic DNA DSB formation could radiosensitize AR-expressing prostate cancer cells with colony-forming potential, VCaP cells were deprived of androgens and stimulated with DHT or FLU for 6 hours, when DSBs are present (**Fig 3A-B**). At 6 hours following exposure to DHT or FLU, cells were exposed to ionizing radiation and then plated at clonal density for clonogenic assay analysis. Pre-treatment of androgen-deprived cells with a 6 hour exposure to flutamide led to a dose-dependent sensitization of VCaP cells to IR, and this sensitization was most notable at 1 and 2Gy of IR (**Fig 4C**). As IR dose increased, radiosensitization decreased, suggesting cell killing is driven by IR at these higher doses, consistent with the observations in synergistic DSB formation (**Fig 4A-B**). To test the ability of FLU to radiosensitize AR-expressing prostate cancer cells *in vivo*, we established VCaP xenografts in surgically castrated athymic, nude mice bearing silastic testosterone implants to control circulating androgen levels. Once palpable tumors formed, silastic implants were removed to mimic androgen deprivation therapy (ADT), and mice were given a subcutaneous injection of sesame oil alone (vehicle) or FLU prior to sham or a tumor-directed single fraction of 6 Gy radiation, and tumor size was monitored with silastic implants replaced to simulate cessation of ADT until tumors grew to 4-fold pre-treatment volume. Untreated tumors grew to 4x pre-treatment volume in approximately 42 days. While single treatments of 6Gy IR or FLU exposure alone led to a tumor growth delay as expected, FLU exposure prior to IR prevented 4x increase in volume for the duration of the study (**Fig 4D**). FLU-induced DNA damage *in vivo* was demonstrated by IHC staining of γH2A.x in biopsy tissues of a cohort of mice bearing VCaP xenografts in parallel with mice being monitored for tumor growth. Biopsies were taken over time and significant increases in γH2A.x foci and nuclear AR were noted at 12 hours, with a return to basal levels by 24 hours, supporting the notion that transient AR-ligand-induced DSBs can radiosensitize AR-expressing tumor cells *in vivo* (**Fig 4E, 4F**).

**Figure 4:**
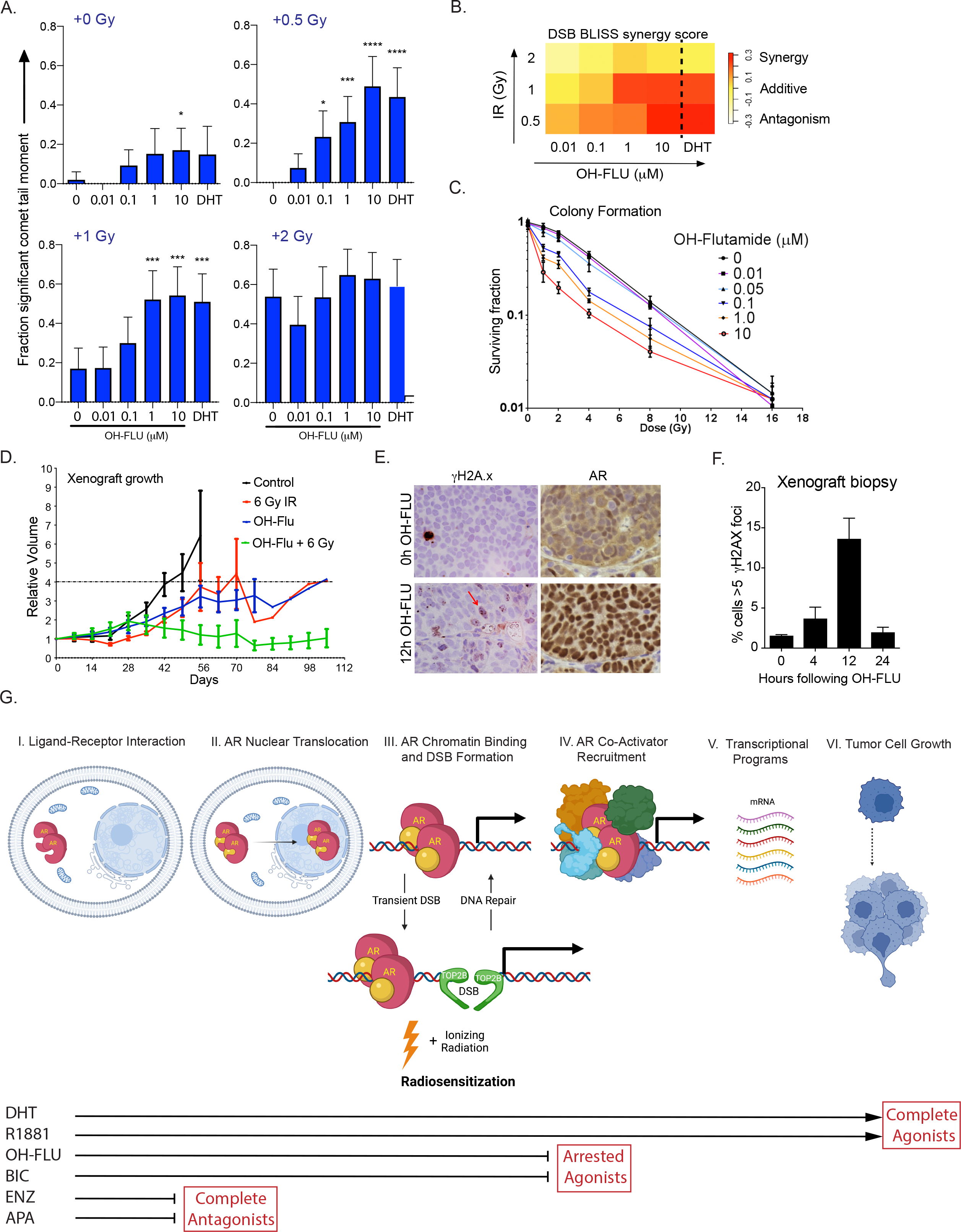
Complete and arrested AR agonist stimulation leads to radiosensitization of prostate cancer cells *in vitro* and *in vivo* due to synergistic double strand break formation. **(A)** Androgen-deprived LAPC4 cells were stimulated with indicated concentration of hydroxyflutamide or 10nM DHT 6 hour prior to 0, 0.5, 1, or 2 Gy ionizing radiation, and immediately placed into the neutral comet assay. Comet tail moment values above 99^th^ percentile are considered significant for at least 50 nucleoids per experiment (*p<0.05, **p<0.01, ****p<0.0001 by 1-way ANOVA and Dunnett post-test compared to vehicle-treated cells). **(B)** BLISS Synergy scores for DSB in combination treatments of hydroxyflutamide (μM) or 10nM DHT prior to indicated dose of IR (Gy). **(C)** Clonogenic potential of androgen-deprived prostate cancer cells following 6 hour exposure to hydroxyflutamide followed by ionizing radiation. Cells were then plated in serum-replete medium at clonal density and colonies are reported as surviving fraction (n=2). **(D)** VCaP xenograft growth. VCaP xenografts were established in surgically castrated mice bearing subcutaneous silastic testosterone-containing implant. Upon reaching approximately 0.1 cm^3^, implants were removed for 16 days prior to stimulation with sesame oil (vehicle) or hydroxyflutamide (50mg/kg) for 12h. Tumors were then radiated at 0 or 6 Gy in a single fraction and tumor volumes were measured over time. **(E)** Representative IHC staining of γH2A.x and AR in biopsies from VCaP xenograft tissues 12 hours following exposure to sesame oil vehicle control (0h OH-FLU) or 50mg/kg hydroxyflutamide (12h OH-FLU). **(F)** γH2A.x foci in biopsies of xenograft tumors were quantified and reported as % cells with >5 foci at 4, 12, and 24 hours following exposure. **(G)** Scheme depicting Arrested Agonist concept exploited to radiosensitize AR-expressing prostate cancer cells. Ionizing radiation-induced DNA damage synergizes with DSB induced by AR complete or arrested agonists. Arrested agonists may provide synergy with radiotherapies while preventing complete AR signaling events.

## Discussion

Here we extend our studies on AR-mediated DNA damage and suggest the concept of *arrested agonism*, wherein some ligands are capable of stimulating a limited cascade of AR signaling steps by virtue of their unique combinatorial assembly of coactivators while inducing little or no canonical target genes or cell growth (**Fig 4G**). Arrested agonism bears key differences from partial agonism, where ligands induce a positive, though limited, agonist effect while competing with agonists. Of note, we provide the first evidence that the BAF complex member SMARCD2 is required for efficient AR agonist-induced signaling and suggest that its absence from the greater nuclear AR complex following OH-FLU exposure in part explains its arrested agonist behavior. BAF complex members have been shown to interact with AR and some individual BAF members such as SMARCE1 have been shown to play a role in AR activity using LNCaP cells^16^. Additionally, recent studies have shown that PROTAC-mediated degradation of the BAF ATPases SMARCA4 and SMARCA2 disrupts super-enhancer and promoter-looping interactions which are involved in oncogenic signaling, highlighting the importance of BAF-mediated nucleosome remodeling in cancer and in prostate cancer specifically^17^. Of note to our study here, the BAF complex has been shown to assemble in an ordered, modular fashion wherein two SMARCC subunits and one SMARCD subunit comprise an initial core heterotrimer upstream of additional core module members followed by an ATPase module which is connected by an Actin-related protein (Arp) module^18,19^. It is possible that the AR influences subunit membership of the BAF complex at an early stage of assembly to include SMARCD2. Interestingly, SMARCD2 was found to interact with the AR following OH-FLU exposure in LNCaP cells, which we speculate is a direct result of the T877A mutation in the ligand binding domain of the AR. OH-FLU has been shown previously and in our study to drive growth of LNCaP cells in castrate conditions as this mutation conveys an antagonist-to-agonist switch in response to various AR antagonists^9-11^, further supporting the idea that SMARCD2 is a critical coactivator in AR signaling and growth in castration sensitive prostate cancer cells.

More broadly, the evidence that an AR antagonist induces DNA DSBs in a manner similar to agonist-induced DSBs and that this damage synergizes with ionizing radiation gives support to the idea of using these molecules in a pulsed schedule paired with radiation to enhance prostate cancer treatment effects. It is unclear whether this feature of AR signaling is unique to the AR. Other hormone-receptor expressing cancers could potentially benefit from pulsed arrested agonist strategies when paired with radiation or other DNA damaging agents. Indeed, hormone induced DNA damage was initially discovered in estrogen receptor-positive breast cancers, though clinical trials did not support estrogen use for this purpose^20,21^. We speculate that a critical feature of proposing hormone receptor-mediated DNA damage with respect to pairing with ionizing radiation would be the pulsatile application of AR ligands to allow for large-scale topological alterations in the genome to occur, which require topoisomerase activity to overcome. While we show that TOP2B is required for agonist and OH-FLU induced DSBs to occur, mechanisms of TOP2B trapping following AR activity have not been elucidated. Overall, our study suggests that arrested agonist-specific combinatorial assembly of AR coactivators could be exploited to radiosensitize AR-expressing prostate cancer cells while preventing full AR signaling-induced cell growth and survival. Moreover, the activity of molecules like OH-FLU herein described may provide a novel pharmacologic concept herein termed “arrested agonism” that could be employed in designing and evaluating small molecules to perform key functions while preventing full agonism, providing opportunities for synergy in co-exposures while simultaneously preventing complete pathway-specific programs.

## Acknowledgements

We thank Jessica Hicks and Muniza Uddin for assistance with IHC staining. John T. Isaacs, Alan K. Meeker, and Karen S. Sfanos provided helpful advice and discussion. Tatiana Boronina and Robert Cole provided expertise in proteomics experiments (Johns Hopkins Mass Spectrometry and Proteomics Core Facility).

## Author contributions

T.L.D and S.Y. conceived the study and all authors contributed to designing experiments. J.B.C., M.C.H., Y.Z., H.Z., M-T.P., R.C., A.M., M.S., S.K., K.C., and A.A. performed the experiments. A.D.M. performed pathology assessments. J.B.C. wrote the manuscript and all authors assisted with the editing the manuscript. All authors contributed to the analysis and interpretation of the data and approved the manuscript in its final form.

## Data and materials availability

All data are available in the main text or the supplementary materials.

## METHODS

### Cell culture and reagents

LNCaP, VCaP, HEK293FT, and CWR22Rv1 cells were purchased from the ATCC (Manassas, VA). LAPC4 cells were obtained from Dr. Charles Sawyers’ laboratory (UCLA, 2002). Cells were grown under conditions previously described and were maintained at low passage numbers. STR genotyping was used to confirm cell line authenticity at 8-12 month intervals. Androgen deprivation was accomplished by washing with 10% charcoal-stripped FBS containing medium twice for 1 hour and incubated with 10% charcoal-stripped FBS for 48-72 hours with daily medium changes prior to the indicated stimulation. Dihydrotestosterone (DHT) was purchased from Sigma-Aldrich. Enzalutamide, hydroxyflutamide, bicalutamide, apalutamide, nilutamide, and darolutamide were purchased from Selleck Chem. Antibodies used were: AR (Millipore-Sigma, 06-680), TOP2B (Novus, NB100-40842), B-Actin (Sigma-Aldrich, A5441), γH2A.x (Millipore, 05-636 clone JBW301), SMARCD2 (Santa-Cruz, 101162), a-Tubulin (Millipore-Sigma, CP06).

### Primers

ChIP primers used were:

KLK3_enhancer F: TGGGACAACTTGCAAACCTG

R:CCAGAGTAGGTCTGTTTTCAATCCA

KLK3_middle F:CAGTGGCCATGAGTTTTGTTTG

R:AACCAATCCAACTGCATTATACACA

KLK3_promoter F: CCTAGATGAAGTCTCCATGAGCTACA

R:GGGAGGGAGAGCTAGCACTTG

TMPRSS2_enhancer F: TGGTCCTGGATGATAAAAAAAGTTT

R: GACATACGCCCCACAACAGA

TMPRSS2_promoter F: CTGAGCCCCCACAATTGC

R: GGTGGGACACACCTCAGCC

RT-PCR primers used were:

KLK3: F: TGAACCAGAGGAGTTCTTGAC R: TGACGTGATACCTTGAAGCA

PRKDC: F: AGCTGGCTTGCGCCTATTT R: GGGCACACCACTTTAACAAGA

GAPDH: F: GGAGCGAGATCCCTCCAAAAT, R: GGCTGTTGTCATACTTCTCATGG

### Immunofluorescence Microscopy

Cells were grown in specified conditions of androgen deprivation or repletion, washed in PBS, and fixed in 3.7% formaldehyde prior to permeabilizing with 0.3% Triton-X100. Primary antibodies were detected using secondaries conjugated with Alexa Fluor 568 or 488 and mounted with DAPI containing mounting medium (Prolong Gold, Thermo Fisher). γH2A.x foci colocalizing with DAPI were quantified using ImageJ (NIH, Bethesda, MD) and the Telometer plugin (JHU) for at least 50 cells per replicate. Nuclear AR levels were quantified using ImageJ (NIH) and reported as pixels per DAPI-positive nuclei. For immunohistochemical labeling of γH2A.X and AR, slides were deparaffinized and rehydrated prior to steaming in HTTR buffer (Dako). Primary antibodies were applied at 1:4,000 (γH2A.X) or 1:300 (AR) dilution at room temperature for 1 h. Immunocomplexes were visualized using PowerVison + Poly HRP from ImmunoVision Technologies Co. using DAB as the chromogen.

### RNAi

shRNA: SMARCD2 knockdown was achieved using the pLK0.1 lentivirus system (Horizon). 2 sequences targeting SMARCD2 (TRCN0000021264, TRCN0000021267) or an empty vector were used to generate lentiviral particles according to published protocols (Broad Institute). Target cells were transduced using polybrene and 48 hours later selected with 5ug/mL puromycin prior to seeding in downstream experiments.

siRNA:3 × 10^5^ LNCaP, VCaP, or LAPC4 cells were seeded per well in their typical growth medium supplemented with 10% FBS. Sixteen hours following plating, siRNAs targeting TOP2B (Hs_TOP2B_6 FlexiTube; Qiagen), AR (SMARTpool: ON-TARGETplus; Dharmacon), SMARCD2 (SMARTpool: ON-TARGETplus; Dharmacon) or nontargeting control siRNA (SMARTpool; Dharmacon) were transfected using RNAiMax transfection reagent (Thermo Fisher). The following day, cells were washed and medium was replaced with growth medium containing 5% charcoal-stripped FBS. Seventy-two hours after transfection, cells in androgen-depleted conditions were exposed to DHT, hydroxyflutamide, or vehicle control. Knockdown was confirmed by Western blotting.

### RIME

Rapid Immunoprecipitation Mass Spectrometry from Endogenous proteins was performed as described previously^22^. Modified from: Mohammed H, et al. Cell Rep. 2013 Feb 21;3(2):342-9. PMID: 23403292. Briefly, LAPC4 cells were grown in steady-state conditions, deprived of androgens for 72 hours in 10% CS-FBS or stimulated with 10nM DHT, 10μM hydroxyflutamide for 3 or 16 hours and nuclear fractions were sonicated using a waterbath sonicator at 4°C (Covaris). Supernatants were precleared using Protein G DynaBeads (Thermo) for 1 hour at 4°C. Pre-cleared lysates were incubated with rabbit anti-AR (Millipore-Sigma, 06-680) or rabbit IgG, followed by incubation with Protein G Dynabeads for 4°C. Immune complexes were collected by magnetization and washed 5 times. On-bead trypsin digest was performed overnight at 37°C. Finally, beads were separated via magnetization and supernatants were collected for MS-analysis.

### Mass Spectrometry

Two replicate samples per condition following IgG or AR immunoprecipitation were submitted for analysis by the JHU Mass Spectrometry Core Facility. Peptides in 40ul 25mM TEAB, the pH ~< 8.0. Protein from IP were digested and the estimated amount from MSMS ID analysis was 10ug. No reduction, alkylation, or other modifications were made to the samples prior to submission for analysis via conventional mass spectrometry protein ID or Isobaric tandem mass tag analysis.

### Peptides labeling with Isobaric mass tags TMT16Pro reagents

70% labels in 14ul acetonitrile used to label peptides in 70uL 100mM TEAB according to Thermo protocol. TMT 16 plex pro-reagent (Thermo Fisher-Pierce, LOT # UH290430. 5 µL of 5% hydroxylamine was added to each sample, then incubated for 15 minutes to quench the reaction. The TMT labeled peptide from each sample was combined into a total 160 000μg combined TMT-labeled peptides, aliquoted 80ug x2, and dried.

### Peptide fractionation by BRP chromatography

80ug of combined TMT labeled peptides, dry, was re-constituted in 50ul 50mM TEAB buffer and cleaned on Pierce Detergent removal columns 0.5mL, to remove excess TMT label, small molecules and lipids. Peptides were eluted in 50uL TEAB. 1950uL bRP buffer A was added followed by 10 mM TEAB in water and peptides were fractionated by basic reverse phase (bRP) chromatography: 2mL injection over 8 min at 250ul/min and fractionated using 85min gradient from 100% solvent A (10mM TEAB in water) to 100% solvent B (90% acetonitrile/10mM TEAB)at flow rate 250ul/min on a XBridge C18 Column, 5 µm, 2.1 x 100 mm column (Waters) with a XBridge C18 Guard Column, 5 µm, 2.1 x 10 mm (Waters), using an Agilent HPLC system 1200 series binary capillary pumps, with variable wavelength UV detector and a micro-fraction collector. 84 fractions were collected and re-combined into 24 fractions for LC-MS/MS analysis.

### LC-MS/MS analysis

Peptides in 24 fractions with calculated average amount per fraction 3.33 µg were re-constituted in 50 µL 2% acetonitrile/0.1% formic acid and 8 µL or calculated average amount / fraction 533 ng analyzed by nano-LC-MS/MS. Peptides were analyzed on a nano-LC-Orbitrap-Lumos-ETD in FTFT (Thermo Fisher Scientific) interfaced with an EasyLC1200 series using reverse-phase chromatography (2%–90% acetonitrile/0.1% formic acid gradient over 78 min at 300 nl/min) on a 75 µm x 150 mm ProntoSIL-120-5-C18 H column 3 µm, 120 Å (BISCHOFF). Eluted peptides were sprayed into an Orbitrap-Lumos-Fusion mass spectrometer through a 1µm emitter tip (New Objective) at 2.4 kV. Survey scans (full ms) were acquired on an Orbi-trap within 375-1600 Da m/z using a Data dependent Top 15 method with dynamic exclusion of 15 s. Precursor ions were individually isolated with 0.7 Da, fragmented (MS/MS) using an HCD activation collision energy 38. Precursor and fragment ions were analyzed at a resolution of 120,000 and 60,000, respectively.

### MSMS data analysis

Tandem MS2 spectra (signal/noise >2) were processed by Proteome Discoverer (v2.4 ThermoFisher Scientific) using Files RC option (recalibration with appropriate database). MS/MS spectra were searched with Mascot v.2.6.2 (Matrix Science, London, UK) against RefSeq2017_83 Human and Rabbit database, and a small database containing enzymes, BSA. Trypsin as an enzyme, missed cleavage 2, precursor mass tolerance 5ppm, fragment mass tolerance 0.01Da and TMT 16pro on N-terminus and TMT 16pro on K as fixed, oxidation on M, deamidation on NQ as variable modifications. Peptide identifications from the Mascot searches were processed within the Proteome Discoverer and Percolator to identify peptides with a confidence threshold of 1% False Discovery Rate, based on an auto-concatenated decoy database search, and calculate the protein and peptide ratios. Only peptide Rank 1 were considered. Only non-modified peptides were used for normalization and ratio calculation. Only Unique peptides were used for ratios calculation. Reporter SN >5, Isolation Interference <25 Proteins appeared in the PD2.4 file were identified at high, medium, and low confidence with at least 1 peptide identified at 1% FDR, Rank1 (Percolator FDR). MSnbase^23,24^ and qPLEXanalyzer^25^ packages were used to organize and analyze protein data from RIME experiments. MS-ARC plot was generated using published methods^22^.

### Cell growth assays

LAPC4 cells transduced with an empty vector (control) or 1 of 2 constructs targeting SMARCD2 were plated in 48-well plates in various conditions: steady state IMDM supplemented with 10% FBS and 1nM R1881 or androgen deprived (10% CS-FBS) and stimulated with 1nM or 10nM DHT. Cell confluence was measured using the incucyte (Sartorius) by taking 16 images of each well every 6 hours over 10 days. Cells in each condition were plated in quadruplicate. Averages of confluence were plotted as percent initial confluence for each condition.

### Coimmunoprecipitation

LAPC4 cells were grown in IMDM supplemented with 10% FBS and 1nM R1881 (steady state). Cells were androgen deprived over 72 hours using IMDM supplemented with 10% CS-FBS and stimulated with 10nM DHT, 10μM hydroxyflutamide, or vehicle (CS). 16 hours following exposure, cells were harvested and lysed in ice-cold RIPA containing 1x protease and phosphatase inhibitors and briefly sonicated. Lysates were incubated with non-immune IgG, anti-AR, or anti-SMARCD2 antibodies overnight followed by 4 hours incubation with magnetic protein G dynabeads (Thermo). Immune complexes were harvested via magnetization, washed 4x with ice cold RIPA buffer, and prepared for western blotting.

### PLA

Proximity ligation assays were conducted using the Duolink Proximity Ligation Assay reagents (Millipore Sigma) according to manufacturer supplied protocol. In brief, cells were on coverslips grown under indicated conditions and then fixed for 10 minutes with 3.7% formaldehyde, washed 3x with PBS, and permeabilized with 0.3% Triton X-100 prior to Duolink PLA assay steps. Images were acquired using a fluorescence microscope (Carl Zeiss AG) and foci colocalizing with DAPI were quantified using ImageJ (NIH, Bethesda, MD) and the Telometer plugin (JHU) for at least 100 nuclei replicate.

### Comet assay

Comet assay experiments were performed as described previously (Hedayati). Neutral pH conditions were used for double strand break detection using a commercial protocol (Trevigen). DNA was detected using SYBR Green and nucleoids were imaged by Zeiss Imager.Z1 fluorescence microscope (Carl Zeiss AG) and analyzed using CometScore (Autocomet.com) software.

### Clonogenic survival

Clonogenic survival was performed as reported previously^13^. In brief, androgen-deprived cells were exposed to the indicated concentration of hydroxyflutamide or vehicle control for 6 hours and then exposed to the indicated dose of ionizing radiation using a MultiRad160 (Precision X-Ray). Cells were seeded in triplicate at clonal density in normal growth medium supplemented with 10% FBS and cells were cultured for 4-6 weeks. Resulting cell colonies were then stained with a solution of crystal violet in 50% methanol and colonies consisting of ≥30 cells were counted.

### Xenograft experiments

All animal experiments were performed in accordance with protocols approved by the Animal Care and Use Committee at Johns Hopkins University. Athymic male nude mice (nu/nu, 8 weeks old) were obtained from the Animal Center Isolation Facility at Johns Hopkins University and maintained in a sterile environment. Mice were castrated and 1-cm-long polydimethylsiloxane (Silastic) implants packed with testosterone (Sigma-Aldrich) were implanted subcutaneously as described previously (8). VCaP cells (3 × 10^6^ cells in 50% Matrigel, 50% PBS) were then injected subcutaneously in the hindlimb. When tumor volume reached approximately 0.1 cm^3^, silastic implants were removed and mice were kept at castrate androgen levels for 16 days. For tumor growth delay studies, mice were treated with one dose of sesame oil (vehicle control) or hydroxyflutamide in sesame oil (50mg/kg) with or without a single fraction of radiation 12 hours following control or hydroxyflutamide exposure. Silastic implants were reimplanted 4 days following treatment and tumor volume was measured using calipers. All xenograft radiation experiments were performed with a single dose of 6 Gy (JL Shepherd Mark 137Cs irradiator; JL Shepherd & Associates). The control group received the same regimen of testosterone withdrawal and replacement, but did not receive any radiation. Tumors were measured every other day to calculate tumor volume (width × length × height × 0.52). Time to 4x tumor growth was determined as the time required for the tumor to reach four times its volume before radiation treatment.

Using an independent cohort of mice for pharmacodynamic studies, castrated VCaP tumor-bearing animals were injected subcutaneously with 50mg/kg hydroxyflutamide dissolved in sesame oil. Serial biopsies were obtained from established tumors immediately before as well as 4, 12, and 24 hours after exposure using an automatic biopsy system (18G x 9 cm; Achieve, Ref A189; CareFusion). Biopsy specimens were then fixed in formalin for 24 hours prior to storage in PBS at 4°C and later embedded in paraffin for sectioning and immunohistochemical evaluation for AR (Millipore-Sigma, 06-680) or γH2A.x (Millipore, 05-636 clone JBW301).

### Graphical representations

Graphical representations of proposed pathways were created with license using BioRender (BioRender.com).

