## Supplemental Figures for "Arrested Agonist Paradigm For Selective Radiosensitization of Prostate Cancer"

Supp. Fig. 1

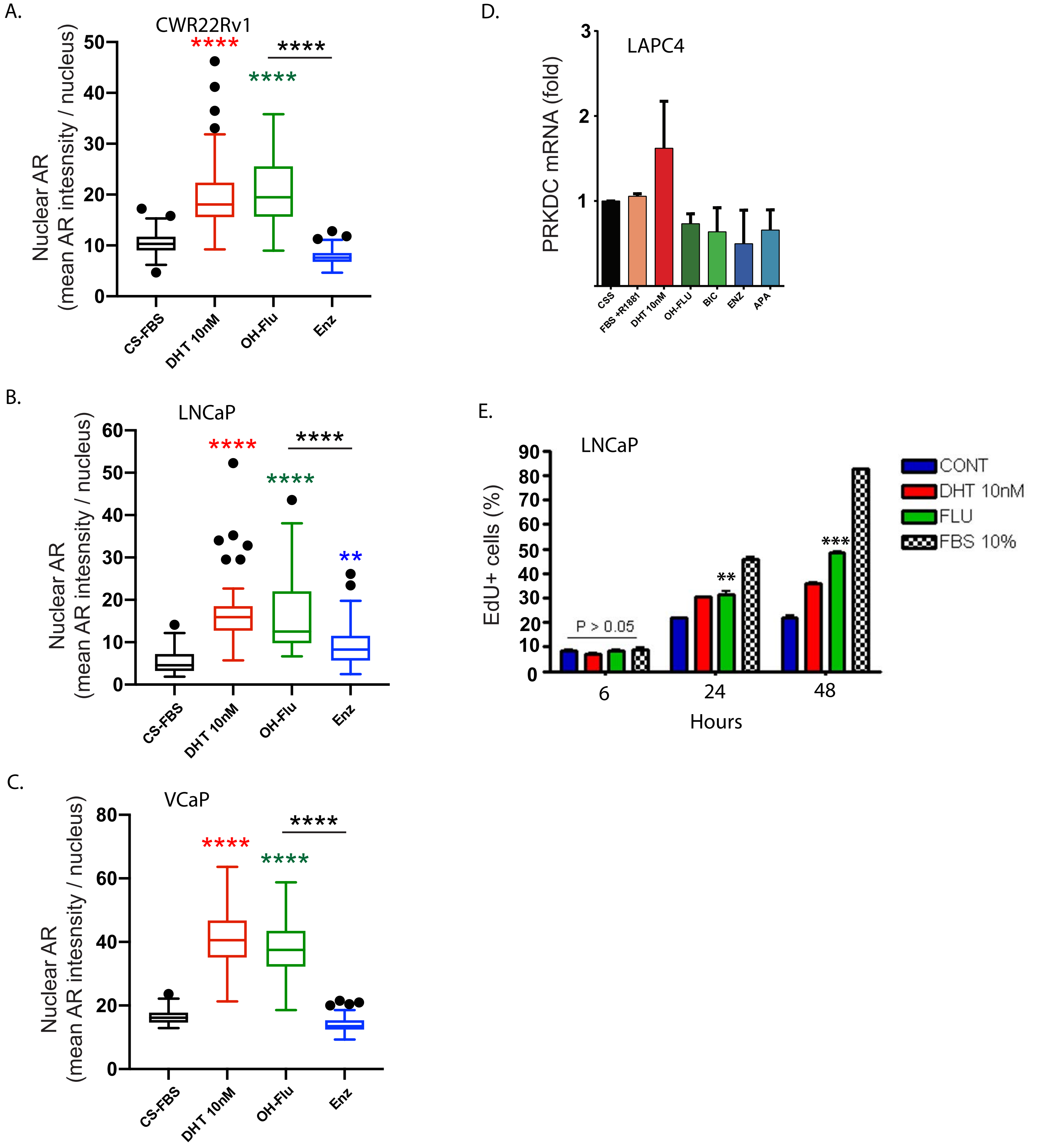


**Supplemental Figure 1: A-C** Immunofluorescent detection of androgen receptor colocalization with DAPI. CWR22Rv1, LNCaP, and VCaP cells were deprived of andogens in CS-FBS medium and stimulated for 2 hours with 10nM DHT or 10μM hydroxyflutamide or enzalutamide. One-way ANOVA with Dunnett's post-test was used to determine significance compared to CS-FBS condition; **p<0.01, ***p<0.001, ****p<0.0001. **D.** PRCDC mRNA following treatment of androgen deprived LAPC4 cells with indicated AR agonist or antagonist. No significance was noted. **E.** DNA synthesis following treatment of androgen deprived LNCaP cells with 10nM DHT, 10uM hydroxyflutamide, or 10% FBS containing growth medium for the indicated time.

Supp. Fig. 2: LRT for individual AR antagonists (FC>0.5)

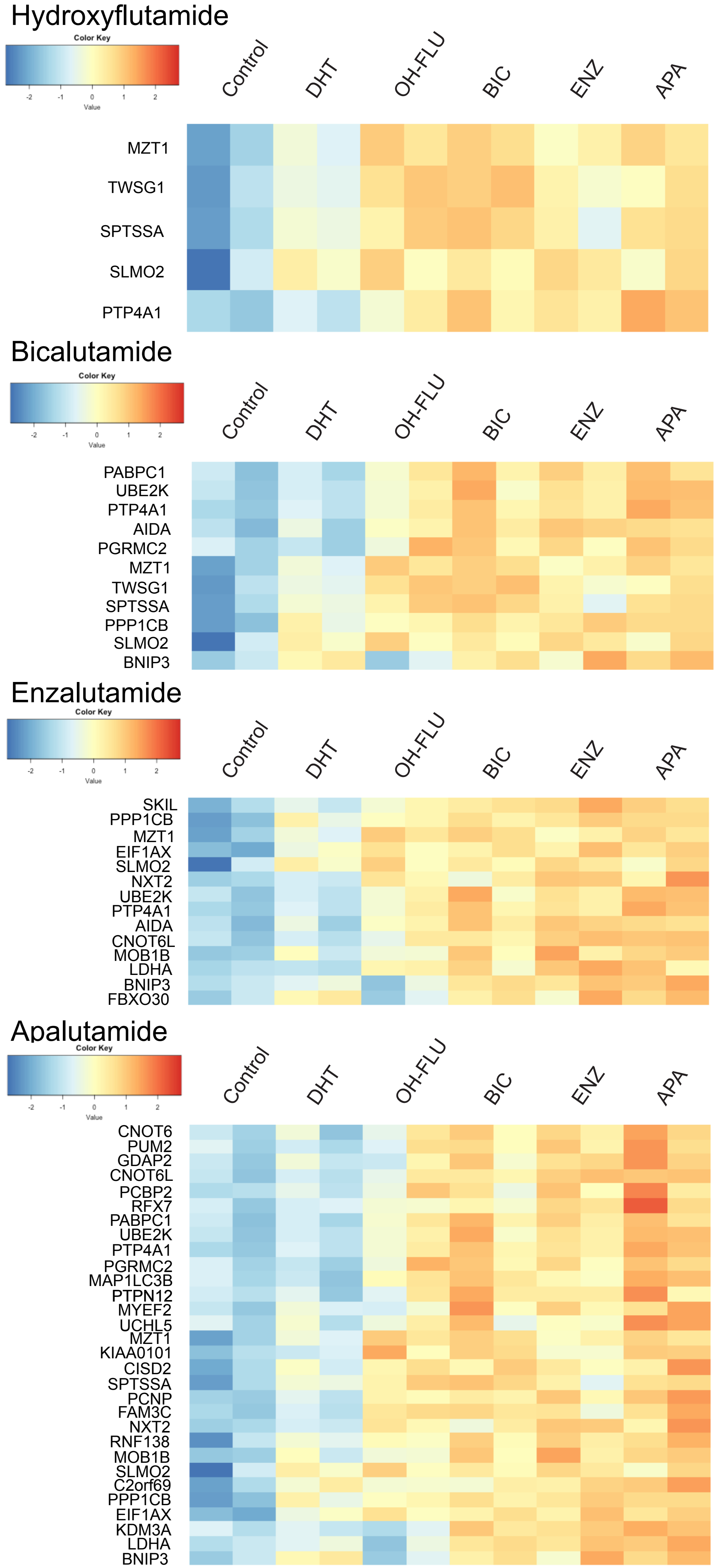


**Supplemental Figure 2:** RNAseq results from LAPC4 cells showing changes in mRNA induced by specific androgen receptor antagonists in the background of androgen deprivation. mRNA targets are top genes induced by the indicated AR antagonist with Log2 fold change >0.5. Samples were run in duplicate for each condition.

Supp. Fig. 3:  Growth curves for individual AR antagonists

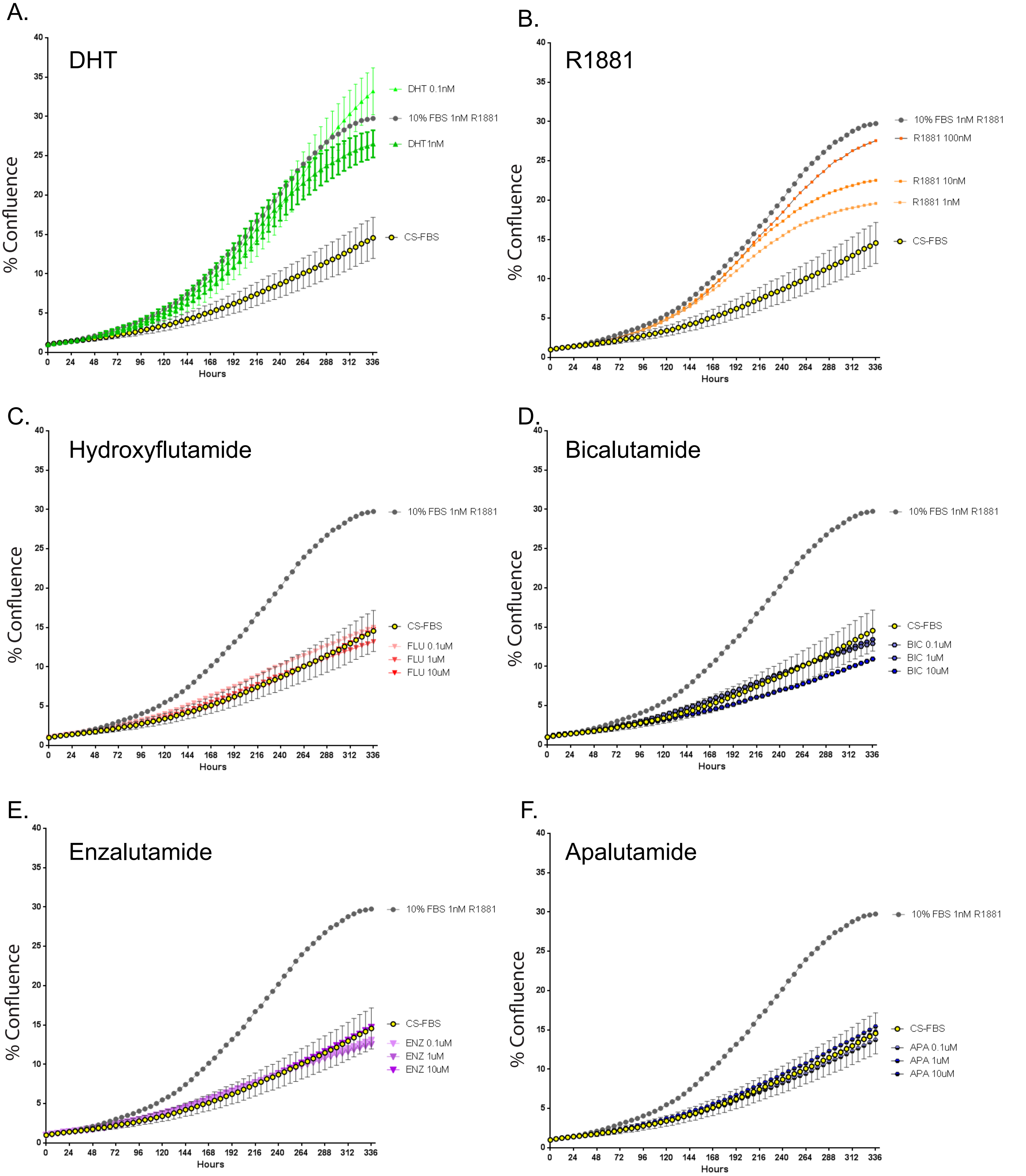


**Supplemental Figure 3:** LAPC4 growth measured by incucyte under androgen deprived conditions (CS-FBS) alone, stimulated with AR agonist (DHT, R1881 alone, or 10%FBS with 1nM R1881), or stimulated with AR antagonist at the indicated dose. Abbreviations: FLU, hydroxyflutamide; BIC, bicalutamide; ENZ, enzalutamide; APA, apalutamide.

Supplemental Fig. 4: Effects of SMARCD2 knockdown

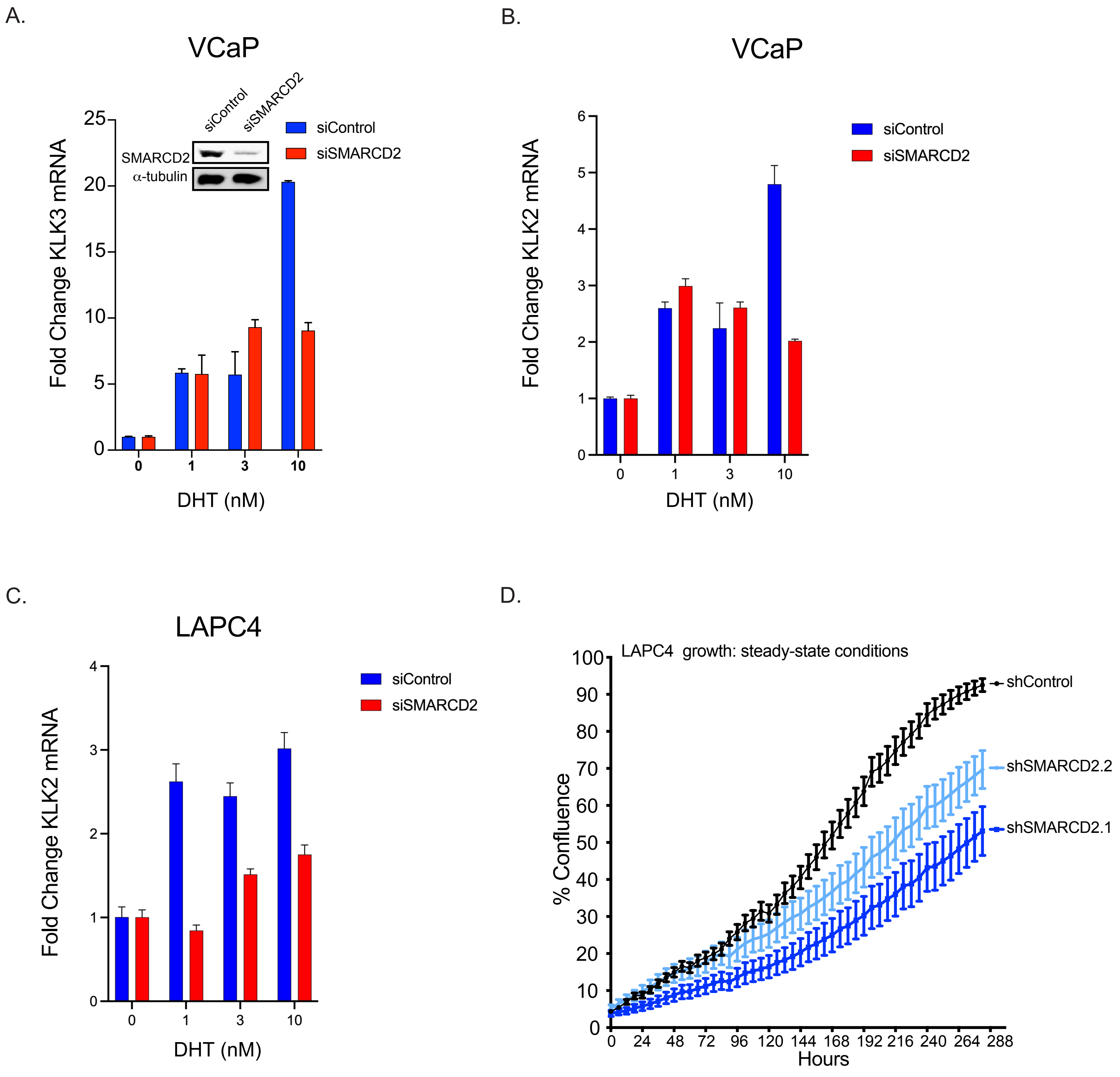


**Supplemental Figure 4:** VCaP or LAPC4 Cells were grown in androgen deprived conditions following knockdown of SMARCD2 or a non-targeted control sequence and then stimulated with the indicated dose of DHT for 24 hours and KLK3 (**A**) or KLK2 (**B, C**) mRNA was measured using GAPDH as a loading control. **D.** Growth of control (shEV) or SMARCD2 knockdown (shSMARCD2.1 or shSMARCD2.2) LAPC4 cells in steady-state, androgen replete conditions.
